# Mutation Impact on mRNA Versus Protein Expression across Human Cancers

**DOI:** 10.1101/2023.11.13.566942

**Authors:** Yuqi Liu, Abdulkadir Elmas, Kuan-lin Huang

**Affiliations:** Center for Transformative Disease Modeling, Department of Genetics and Genomic Sciences, Tisch Cancer Institute, Icahn Institute for Data Science and Genomic Technology, Icahn School of Medicine at Mount Sinai, New York, NY 10029 USA

**Author notes:** Corresponding Author: Kuan-lin Huang, Ph.D., Department of Genetics and Genomic Sciences, Icahn School of Medicine at Mount Sinai, New York, NY 10029.

## Abstract

Cancer mutations are often assumed to alter proteins, thus promoting tumorigenesis. However, how mutations affect protein expression has rarely been systematically investigated. We conduct a comprehensive analysis of mutation impacts on mRNA- and protein-level expressions of 953 cancer cases with paired genomics and global proteomic profiling across six cancer types. Protein-level impacts are validated for 47.2% of the somatic expression quantitative trait loci (seQTLs), including mutations from likely “long-tail” driver genes. Devising a statistical pipeline for identifying somatic protein-specific QTLs (spsQTLs), we reveal several gene mutations, including *NF1* and *MAP2K4* truncations and *TP53* missenses showing disproportional influence on protein abundance not readily explained by transcriptomics. Cross-validating with data from massively parallel assays of variant effects (MAVE), *TP53* missenses associated with high tumor TP53 proteins were experimentally confirmed as functional. Our study demonstrates the importance of considering protein-level expression to validate mutation impacts and identify functional genes and mutations.

## INTRODUCTION

Cancer arises from the acquisition of mutations that confer selective advantages. The majority of these mutations are thought to affect cellular functions by regulating the expression of gene products. For example, truncations can result in nonsense-mediated decay (NMD)^1,2^, which protects eukaryotic cells through degrading premature termination codon (PTC) bearing mRNA^3^. Additionally, a fraction of cancer mutations may uniquely affect protein abundance but not mRNA expression. However, previous studies characterizing genomic mutations affecting mRNA vs. protein levels have focused on germline variants as expression quantitative trait loci (eQTL)^4–6^. While other cancer studies have characterized the effect of somatic mutations on mRNA expression levels^7–9^, it remains unclear how somatic mutations may affect protein abundance. The gap of knowledge is critical given that mRNA and protein levels are only moderately correlated^10–13^. A myriad of factors, including cell state transition, signal delay, translation on demand, and cellular energy constraint, can lead to discrepancies between mRNA and protein levels^14^. Understanding protein-level consequences of cancer mutations is critical in identifying functionally important mutations and revealing their downstream mechanisms.

In recent years, advances in mass spectrometry (MS) technologies have generated a wealth of global proteomic profiles of primary tumor cohorts, many of which also have concurrent genomic and transcriptomic profiling^15–20^. These proteogenomic datasets present ample opportunities to validate somatic mutations that show concordant impacts on downstream mRNA and protein levels. On the other hand, protein abundance may also be uniquely influenced by the efficiency of protein translation efficiency, transport, and degradation. Thus, proteogenomic analyses can reveal mutations that disproportionally impact protein abundances that may not be found using genomic analyses alone.

Herein, we conducted a systematic analysis to decode the relationship between somatic mutations vs. mRNA and protein levels using data from nearly a thousand cases across six cancer types in prospective and retrospective cohorts from the Clinical Proteomic Tumor Analysis Consortium (CPTAC). We identified mutations showing concordant effects at both mRNA and protein expression levels *in cis*, as well as those that showed protein-specific effects. We further examined how mutations associated with expression changes may predict *in vitro* and *in vivo* functional effects measured by a massively parallel assays of variant effects (MAVE) of TP53^21^. Our results highlight the importance of pairing genomic and proteomic analyses to prioritize functionally important mutations.

## RESULTS

### Mutation impacts on the mRNA and protein levels

Following the study workflow (**Figure 1A**), we first sought to identify somatic mutations that may impact the corresponding gene’s mRNA expression (somatic eQTL, termed seQTL below) and protein abundance (somatic pQTL, termed spQTL below) in primary tumor tissue samples. We performed a multiple regression analysis adjusted for age, gender, ethnicity, and TMT batch using the prospective CPTAC datasets that included matched DNA-Seq, RNA-Seq, and mass spectrometry (MS) global proteomics data of primary tumor samples across six cancer types (**Methods**, **Figure 1B**), including 115 breast cancer (BRCA)^19^, 95 colorectal cancer (CRC)^16^, 110 clear cell renal cell carcinoma (CCRCC)^15^, 109 lung adenocarcinoma (LUAD)^17^, 84 ovarian cancer (OV)^20^, and 97 uterine corpus endometrial carcinoma (UCEC)^18^, as well as proteogenomic datasets for additional, retrospective BRCA^11^, CRC^13^, and OV^12^ cohorts from CPTAC for validation (**Figure S1A**). We focused on coding mutations given the coverage of the whole-exome sequencing (WES) data used in CPTAC studies; the analyses were further stratified for truncations, missense, and synonymous mutations given their likely different mechanisms of action in affecting levels of the mutated gene product.

**Figure 1.**
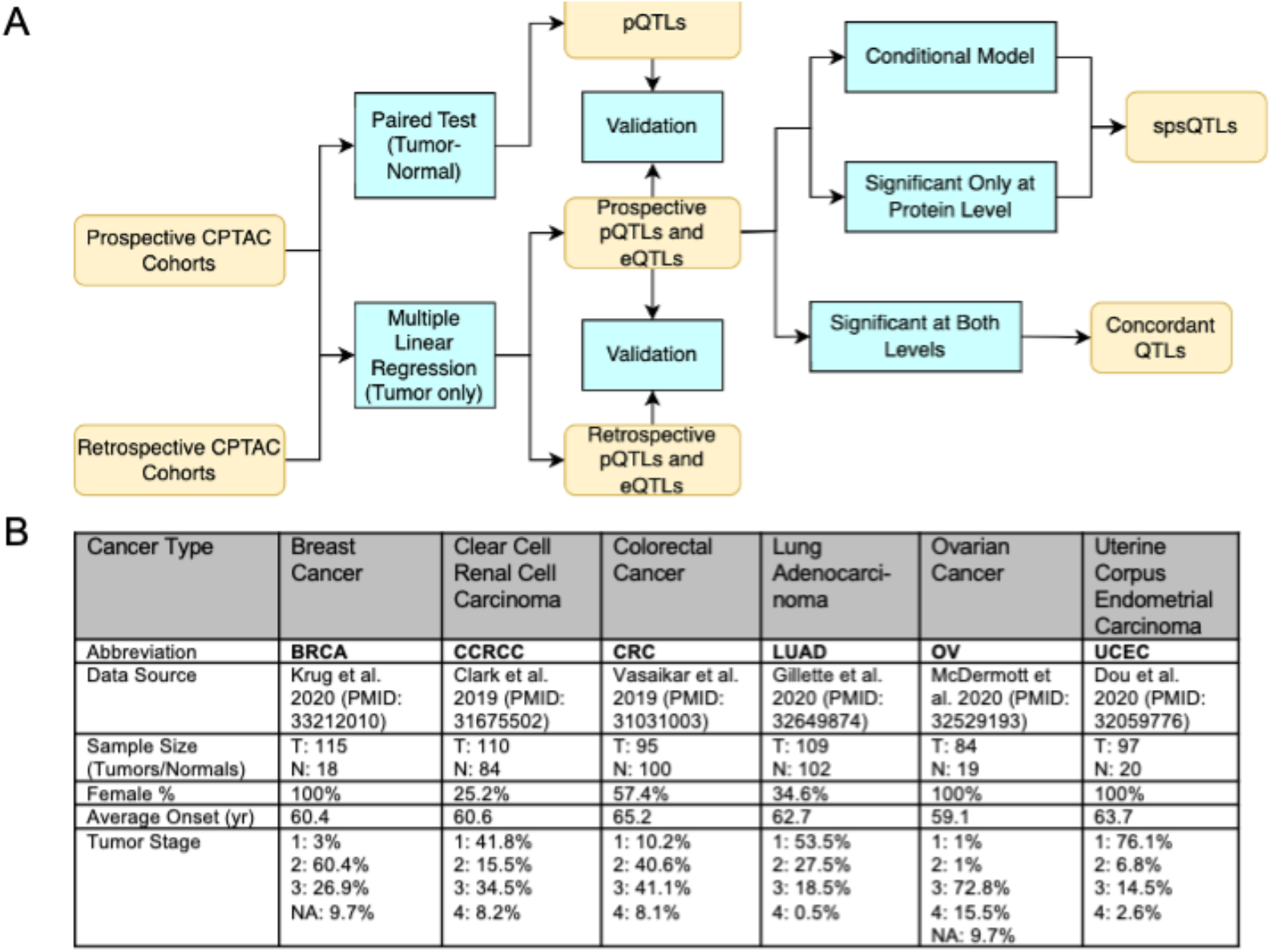
Overview of the proteogenomic cohorts and schematics. (A) Study workflow to identify eQTLs, pQTLs, concordant QTLs (between mRNA and protein levels), and spsQTLs showing disproportional effects on protein expression. (B) Summary of the prospective CPTAC proteogenomic cohorts used for the discovery analyses, including cancer type abbreviation, data source, sample size of tumor (T) and normal (N) tissues, female percentage, average onset age in years, and tumor stage distribution.

Based on the statistical power achieved by these cohort sizes and to reduce false positives, we focused on genes with three or more samples affected by mutations in each functional class of missense, truncation, and synonymous within the cancer cohort, including 134, 13, and 15 genes tested in BRCA; 1360, 318, and 226 genes tested in CRC; 55, 12, and 4 genes tested in CCRCC; 94, 4, and 8 genes tested in LUAD; 134, 5, and 8 genes tested in OV; 2243, 273, and 196 genes tested in UCEC. We sought to identify their seQTLs affecting *cis-*expression, i.e., expression of the mutation-affected genes. Using the multiple regression model (**Methods**), we identified 74 gene-cancer seQTL pairs (FDR < 0.05), including 4 in BRCA, 47 in CRC, 7 in CCRCC, 3 in LUAD, 1 in OV, and 12 in UCEC (**Figure 2A**). Separated by the functional classes of mutations, 22 of those seQTLs are missense mutations, 12 are synonymous, and 40 are truncating. Top seQTLs showing up-regulation of gene expression are primarily missenses, including *SMARCA4* in LUAD, *WNT7B* in CRC, *TP53* in OV, and *FOXR2* in UCEC. Top candidates showing down-regulation of gene expression include *TP53* and *CDH1* truncations in BRCA, as well as *TP53* truncations in OV (**Figure 2B**).

**Figure 2.**
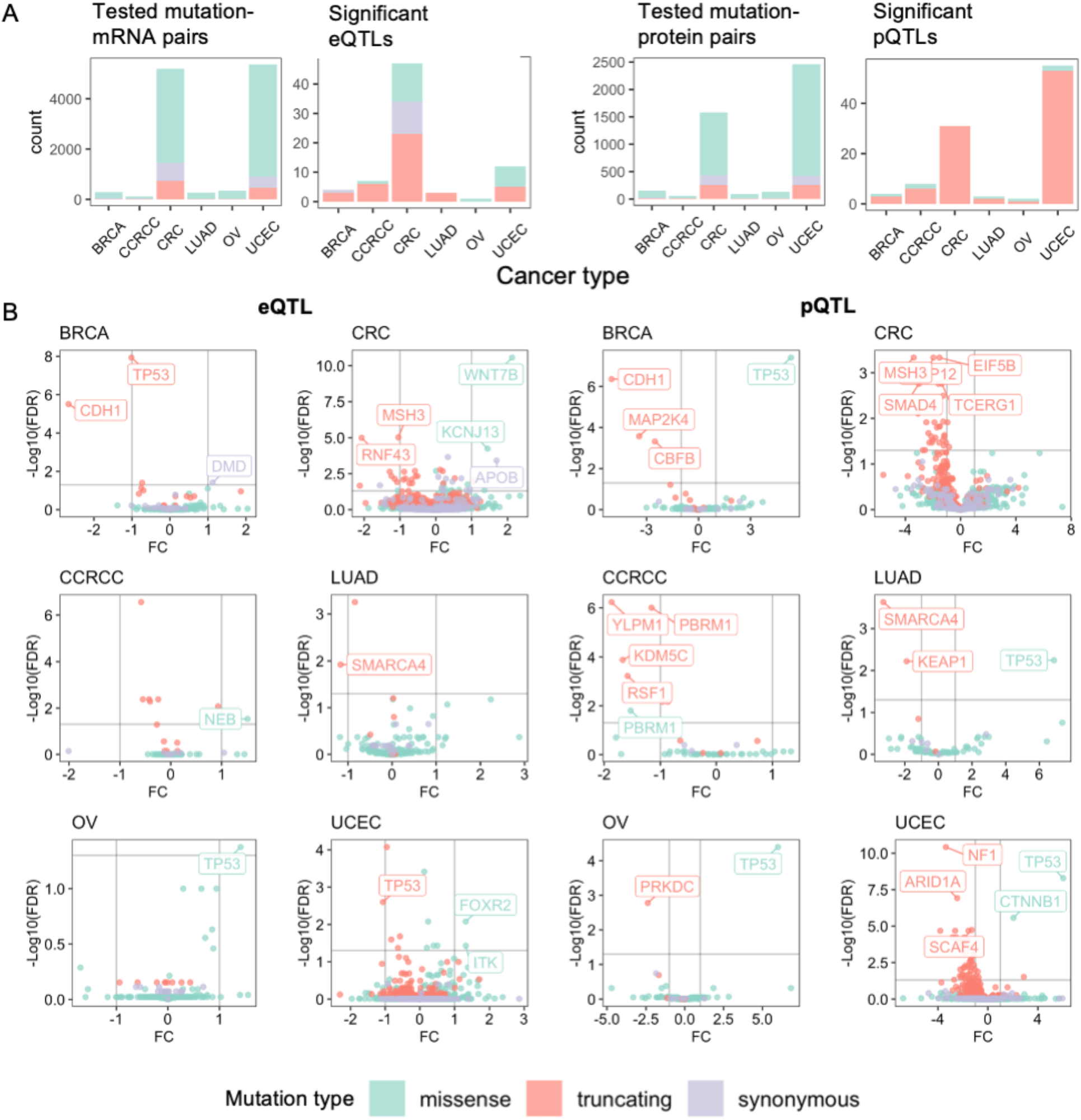
Gene mutations identified as *cis* seQTLs and spQTLs across six adult cancer types. (A) Overview of the somatic mutation QTLs identified in different cancer types and mutation types, including missense (green), truncating (orange), and synonymous (purple) mutations. For both eQTLs and pQTLs, the panel on the left shows the counts of the mutation-gene pairs included in analyses, and the figure on the right shows the counts of the significant eQTLs and pQTLs. (B) Volcano plots showing seQTLs associations in the six cancer types (left) and volcano plots showing spQTLs associations (right), where each dot denotes a gene-cancer pair included in the analysis. Top associated genes were further labeled. FC: mRNA/protein expression log fold change. FDR: false discovery rate.

Using a similar multiple regression but modeling protein abundance as the dependent variable, we identified 103 significant gene-cancer spQTL pairs (FDR < 0.05), including 4 in BRCA, 31 in CRC, 8 in CCRCC, 3 in LUAD, 2 in OV, and 55 in UCEC (**Figure 2A**). Compared to the proportion of gene-mutation type evaluated in each cancer type, spQTLs showed significant enrichment for truncations (Fisher exact test p-value < 0.05; **Figure 2A**), highlighting the persistent and more profound effect of truncations on protein abundance compared to mRNA levels. Among the identified spQTLs across cancer, 7 are missense and 96 are truncating. For example, truncating mutations of *NF1* and *ARID1A* in UCEC, and *YLPM1* in CCRCC are each associated with reduced protein level of the corresponding gene (**Figure 2B**). Notably, *TP53* missenses in OV, BRCA, LUAD, and UCEC are each significantly associated with increased protein expression in mutation carriers (**Figure 2B**).

To verify these discoveries, we applied the same seQTL and spQTL analyses using retrospective CPTAC data (**Figure S1A**) that included independent cohorts of BRCA^11^, CRC^13^, and OV^12^ primary tumors. While these cohorts afforded smaller sample sizes, 8 seQTLs and 5 spQTLs were detected in both retrospective and prospective sets. The gene-cancer spQTL pairs showing strong validation in both datasets include *TP53* missense mutations and *CDH1* truncations in BRCA, and *TP53* truncations in CRC (**Figure S1B**).

### Mutations showing concordant effects at mRNA and protein levels

We next examined the concordance of seQTL and spQTL associations for each gene-cancer type pair. As expected, for most (88.9%) of the significant seQTLs whose genes had sufficient observations at both the mRNA and protein levels, the identified associations showed the same directionality. However, we only identified 17 seQTLs (47.2%) that are also significant spQTLs at an FDR < 0.05, which we show as concordant QTLs (**Figure 3A**). The effect sizes (in log fold change) of these gene-cancer pairs showing concordant seQTLs and spQTLs showed a high correlation between mRNA and protein (Pearson r = 0.90, p-value < 7.51E-7).

**Figure 3.**
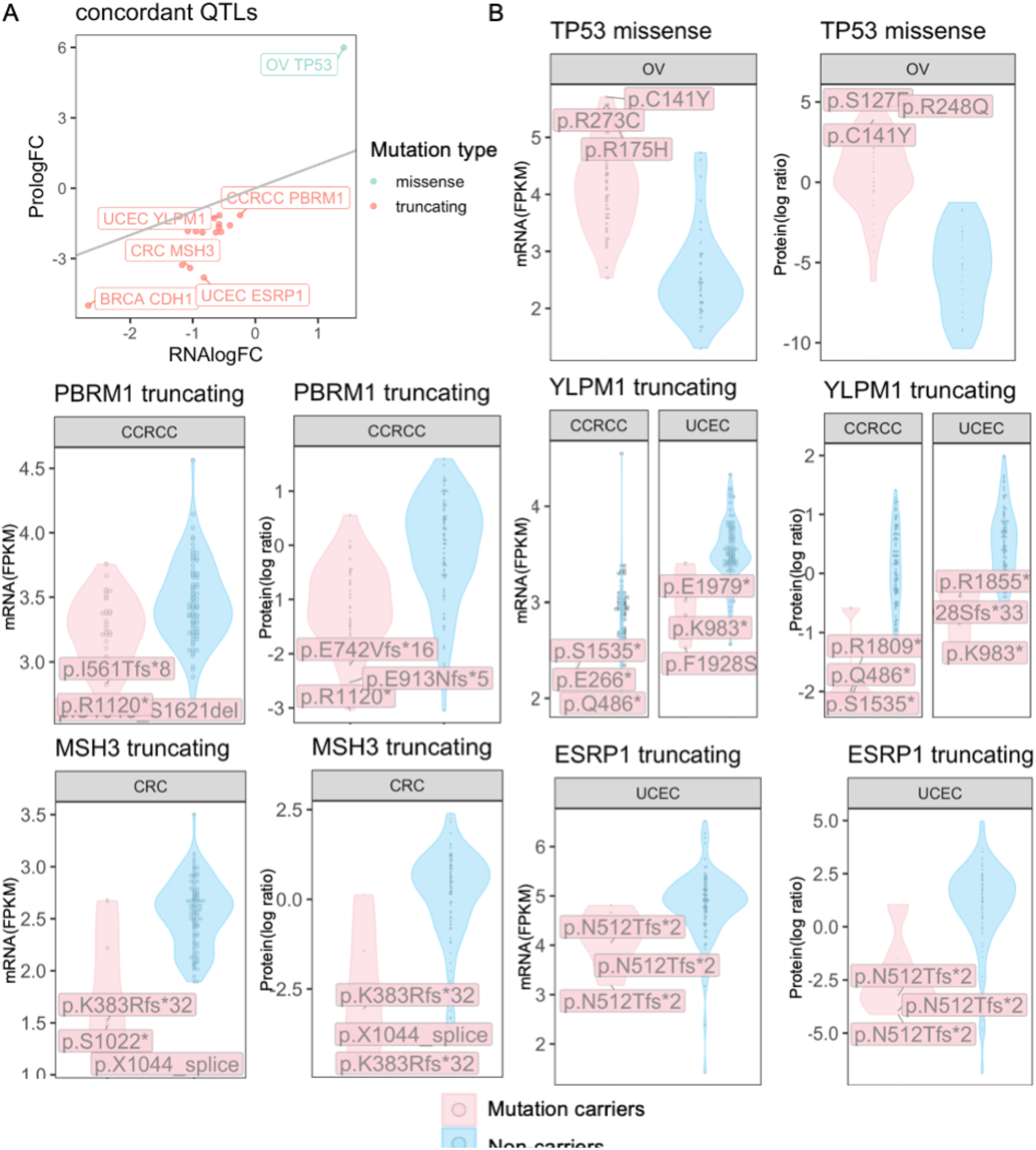
Gene mutations showing concordant impacts on gene and protein expression levels. (A) Overview of concordant QTLs as shown by their effect sizes in log[Fold Change (FC)], where the gray line shows when the protein logFC equals RNA logFC. Some of the top concordant QTLs were further labeled by cancer type and gene name. (B) Examples of QTL with concordant effects at mRNA and protein expression levels. For each gene, the plot on the left shows the corresponding mRNA levels of mutation carriers vs. non-carriers in FPKM, and the plot on the right shows protein level comparison in log ratio (MS TMT measurements) in the respective cancer type labeled on top of each of the violin plots. The labeled mutations are the three mutations whose carriers show the highest absolute expression differences of the mutated gene product compared to the non-carriers.

In different cancer types, genes whose mutation impacts on gene and protein expressions are concordant include well-known drivers of the disease, including *TP53* missense mutations in OV, *CDH1* truncations in BRCA, and *MSH3* truncations in CRC. Up-regulation of mutated *TP53* in OV is the only association found for genes affected by missense mutations. The 16 other concordant se/spQTLs are all truncations associated with reduced expression and highlight some “long-tail” driver genes, including *PBRM1* in CCRCC, *YLPM1* in CCRCC/UCEC, and *ESRP1* in UCEC (**Figure 3B**). The concordant QTLs with truncating mutation can likely be explained by NMD, which reduces gene expression and in turn diminishes the expression of the corresponding proteins^3^. Compared to the substantially higher counts of seQTL associations (**Figure 2A-B**), these concordant se/spQTL effects validate mutation impacts on the gene product.

### Protein-specific mutation impacts not observed at mRNA levels

While most seQTLs and spQTLs show concordance, we postulate that certain mutations may uniquely affect protein abundance but not mRNA levels, which we term somatic protein-specific QTLs (spsQTLs). To identify spsQTLs, we applied two methods to stringently retain QTLs with discordant effects at mRNA and protein levels. First, applying a likelihood ratio test (LRT) between two regression models of protein level being predicted by mRNA level with or without the mutation term (**Methods**)^4^, 96 candidate spsQTLs (FDR < 0.05) were identified. Second, complementing this LRT test with an approach filtering for gene-cancer pair showing significant spQTL (FDR < 0.05) but not seQTLs (**Methods**) ^22^, 86 candidate spsQTLs (FDR < 0.05) were identified.

By overlapping candidate spsQTLs identified by both methods, we retained 83 spsQTLs, the majority (92.8%) of which are truncating mutations (**Figure 4A**). Top spsQTLs associated with diminished protein expression include *NF1* truncations in UCEC, *PLEAHK5* truncations in CRC, and *MAP2K4* truncations in BRCA. The only spsQTLs that increase protein expression include *TP53* missense mutations in BRCA, LUAD, and UCEC. (**Figure 4B**). We further examined the discordance in mutation impacts on gene and protein expression levels (**Figure 4C**). While some of these truncations, such as *NF1* in UCEC and *MAP2K4* in BRCA, were often accompanied by lower-than-median mRNA expression in their respective tumor cohorts, their impacts were strikingly observed at diminished protein expression levels. We highlighted in **Figure S2A** spsQTLs where the affected gene’s protein showed negative protein log fold-change (logFC) whereas the mRNA logFC is non-negative, including *CASP8* truncations in UCEC, *ARID1A* truncations in CRC and UCEC, and ATM truncations in LUAD and UCED. We also identified a set of spsQTLs truncations, where the logFC associated with a reduction in proteins is 15 times greater than mRNAs logFC (**Figure S2B**). These results suggest that NMD associated with these gene truncations are closely tied to the terminated translation but may not affect mRNA expression to the same degree ^23^.

**Figure 4.**
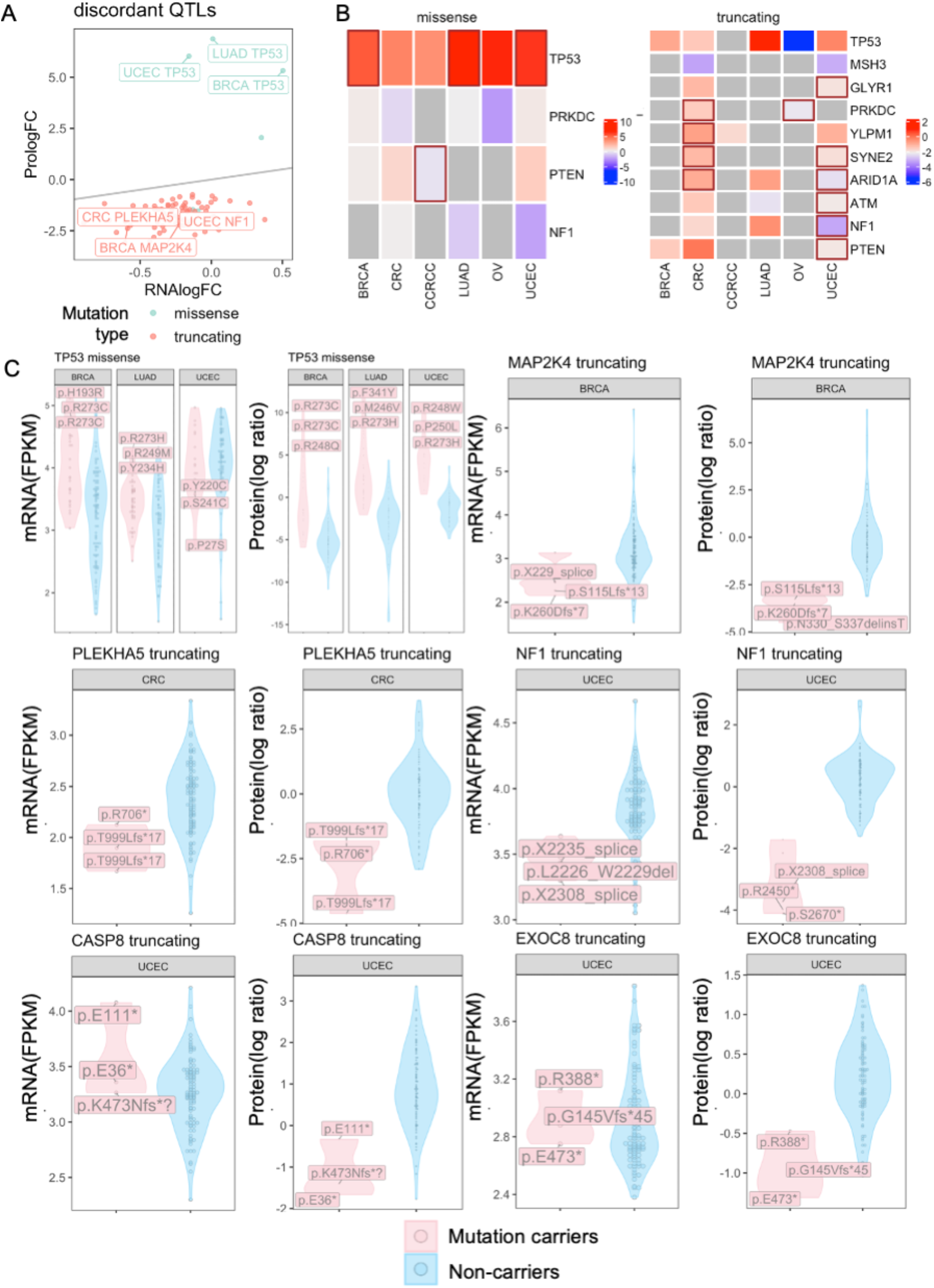
Gene mutations showing discordant impacts on gene and protein expression levels. (A) Overview of discordant QTLs identified by our statistical pipeline as shown by their effect sizes in log[Fold Change (FC)], where the gray line shows when the protein logFC equals RNA logFC. (B) Heatmaps of QTLs that are significant as either seQTL or spQTL and that are shared across at least two cancer types. Brown box indicates significant spsQTLs, and color indicates the effect size in log[Fold Change (FC)], average protein expression of mutation carriers in log ratio from the MS TMT quantifications. (C) Examples of QTL with discordant effects at mRNA vs. protein levels. For each gene, the plot on the left shows the corresponding mRNA levels of mutation carriers vs. non-carriers in FPKM, and the plot on the right shows protein level comparison in log ratio (MS TMT measurements) in the respective cancer type labeled on top of each of the violin plots. The labeled mutations are the three mutations whose carriers show the highest absolute expression differences of the mutated gene product compared to the non-carriers.

To complement the cross-tumor analyses, we also utilized the CPTAC samples with paired tumor-normal tissues to conduct paired differential expression tests for both protein and mRNA expression (**Figure 1A**). The paired sample sizes with proteomic data include 17 in BRCA, 17 in UCEC, 84 in CCRCC, 100 in LUAD, 29 in CRC, and 10 in OV (**Figure 1B**). Covariates including age at diagnosis, ethnicity, race, and sequencing operator are adjusted in the analysis. While this analysis had varied statistical power due to different normal tissue availabilities across cancer types, it served as an independent validation of spQTLs. This paired tumor-normal analysis validated the protein-level impacts of several discordant spsQTLs (**Figure S3A**) as well as some concordant se/spQTLs (**Figure S3B**). For example, the validated discordant spsQTLs include truncations of *SMAD4* and *SCRIB* in CRC as well as *NF1, GLYR1,* and *RASA1* in UCEC (**Figure S3A**). The validated concordant se/spQTLs include truncations of *YLPM1 and PBRM1* in CCRCC, *SMARCA4* and *KEAP1* in LUAD, and *ESRP1* as well as *JAK2* in UCEC (**Figure S3B**).

### Functional evidence of *TP53* missenses associated with high protein expression

Notably, *TP53* missenses are associated with higher protein expression in multiple cancer cohorts, in addition to the expected reduction in expression associated with truncations (**Figure 5A**). Such cis-effect of functional *TP53* missense mutations had previously been observed through immunohistochemistry (IHC^24^) or MS global proteomics experiments^25^. Here, we hypothesized that functional *TP53* missense mutations are more likely to show high levels of concurrent protein-level expression in the mutated tumor sample. To test this hypothesis, we compared gene and protein-level *TP53* expression from CPTAC with *TP53* mutation-level functional data from the *in vitro* and *in vivo* MAVE experiment conducted by Kotler et al^21^, where they designed a p53 variants library to study the functional impact of those mutations.

**Figure 5.**
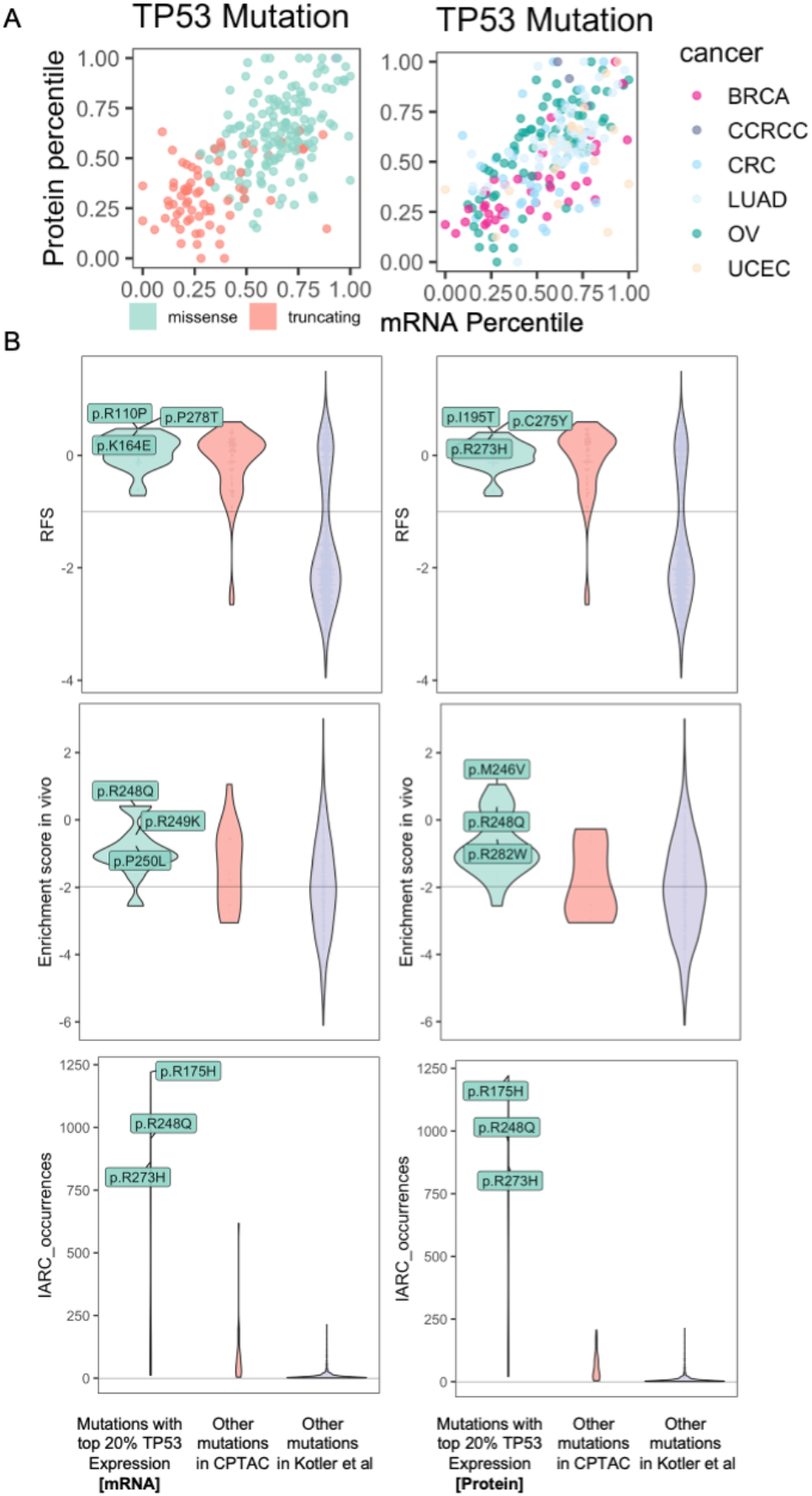
Functional verification of *TP53* mutation associated with high mRNA or protein levels using *in vitro* and *in vivo* data from a MAVE experiment. (A) Percentile of averaged expression associated with a given *TP53* mutation at the mRNA (x-axis) and protein (y-axis) levels in the respective cancer cohort. *TP53* mutations are color coded by mutation type (left) and observed cancer type (right), respectively. (B) Violin plots comparing the in vitro functional score (RFS, top), in vivo enrichment score (middle), and IARC occurrences (bottom) for TP53 mutations in the three groups defined by (1) *TP53* mutations with top 20% mRNA (left) or protein (right) expression in the prospective CPTAC cohorts, (2) the other *TP53* mutations observed across all CPTAC samples, and (3) the rest of the assayed *TP53* mutations from Kotler et al^21^.

We divided the *TP53* missense mutations from Kotler et al. into three categories: (1) *TP53* mutations with top 20% mRNA or protein expression in the prospective CPTAC cohorts, (2) the other *TP53* mutations observed across all CPTAC samples, and (3) the rest of the assayed *TP53* mutations from Kotler et al. For *in vitro* data, the number of tested mutations by each category is 32, 78, and 1,033, respectively. For *in vivo* data, the number of tested mutations by each category is 19, 10, and 381, respectively. We first compared the relative fitness score (RFS) measured from the *in vitro* assays^17^. While there may be a trend, we did not observe a significant difference between all the other mutations versus *TP53* missenses associated with either top 20% expression based on either mRNA (p-value = 0.090, Wilcoxon rank-sum test) or protein expression (p-value = 0.720).

We next compared the *in vivo* enrichment scores across the same categories, and found that *TP53* missenses associated with top 20% protein expression showed significantly higher enrichment score *in vivo* compared to that of other *TP53* missenses found in CPTAC (p-value = 0.016) or other experimentally-measured *TP53* mutations (p-value = 3.23E-5, **Figure 5B, Table S2**). In comparison, *TP53* missenses associated with top 20% mRNA expression did not show a significant *in vivo* score difference to that of other *TP53* missenses found in CPTAC (p-value = 0.170). Kotler et al. observed that there was no significant correlation between enrichment score *in vivo* and RFS *in vitro*, which is consistent with our observations and may be explained by the different selective pressures between these settings *in vivo* and *in vitro*^21^. Finally, *TP53* missenses associated with top 20% protein expression (p-value = 5.91E-7) or top 20% mRNA expression (p-value = 2.38E-2) showed significantly higher prevalence than other CPTAC mutations based on counts from the International Agency for Research on Cancer (IARC) database^21^ (**Figure 5B, Table S2**). Overall, these analyses suggested that protein-level consequences from primary tumor samples can aid the identification of functional mutations.

## DISCUSSION

Herein, we analyzed how somatic mutations affect mRNA and protein levels using matched genomic, transcriptomic, and global proteomic data from 953 cases across six solid cancer types. We first investigated the mutation impacts at the mRNA level and protein level, finding that although most seQTLs have the same direction of effect as spQTLs, less than half of them are also significant at the protein level. We also studied the concordant or discordant relationship between seQTL versus spQTLs, finding several spsQTLs that have disproportional effects on protein. Finally, we conducted analyses to provide functional validation^21^ for our findings of TP53 missenses associated with high protein expression.

Integrating protein-level data identified nearly 47.2% seQTLs as concordant, significant spQTLs. The result demonstrates the capacity of proteomic data to validate genomic findings and potentially filter out noises that may arise for example due to the more transient nature of transcription compared to translation. In addition to well-known tumor suppressors like *TP53* and *MSH3*, other gene mutations with concordant effects may also be “long tail” driver genes that will otherwise require large cohort sample sizes to discover. For example, *PBRM1*, which we found in CCRCC, is a subunit of the PBAF chromatin remodeling complex thought to be a tumor suppressor gene whose mutations may confer synthetic lethality to DNA repair inhibitors^26^. *ESRP1*, found in UCEC, is crucial in regulating alternative splicing and the translation of some genes during organogenesis^27^. Other less-studied genes we identified include *YLPM1* truncations associated with concordantly reduced *YLPM1* mRNA and protein expression levels in both CCRCC and UCEC. Analyzing the distribution of these gene mutations on NCI’s Genome Data Commons, we observed many other recurrent truncations (**Figure S5**), suggesting these mutations may represent some of the “long tail” driver mutations that warrant further investigation^28,29^.

By devising a specific pipeline to detect spsQTLs, our results showed that apart from mutations that influence protein level mediated by changes in mRNA level, many mutations are associated with disproportional aberrations at the protein level compared to mRNA changes, indicating post-transcriptional regulation. SpsQTLs were found to affect known driver genes such as *TP53* missenses, and truncations in *NF1*^30^ and *MAP2K4*^31^. In most cases, protein molecules are more direct mediators of cellular functions and phenotypes than mRNAs^32^. Thus, the discordant effect between mRNA level and protein level discovered in our study highlights the importance of exploring disease mechanisms and developing treatments at the protein level.

This study has several limitations. First, our findings do not distinguish between several potential mechanisms that could lead to discordant effects of mutations on gene and protein expression. One possibility is that the mutation affects the efficiency of translation, leading to changes in protein levels that are not reflected in mRNA levels. For example, accumulating evidence in recent years suggests that NMD is closely tied to the termination of translation^23^, which may explain instances where some truncations afford much stronger associations with protein levels. The mechanisms of how mutations may affect protein translation may be context- and gene-specific and remain to be elucidated. Second, the proteogenomic tumor cohorts used herein, while being some of the largest studies to date, still are limited in sample sizes and preclude sufficient statistical power to identify pQTLs at a single mutation level or reveal *trans* effects. Third, given the limitation of current omic technology and data, our findings do not resolve mutation impact on proteins at the temporal, spatial, or single-cell resolution, but provide candidate mutations to be investigated in future studies.

Finally, using *TP53* missense mutations as an example, we showed that protein-level expression can serve as an effective strategy to prioritize functional mutations. As DNA-Seq become ever more commonplace, many rare mutations are being identified and it remains challenging to accurately classify their functional impacts. Our data demonstrated that *TP53* missenses associated with high protein expression show significantly higher functional scores, particularly those measured *in vivo*. This protein-expression-based prioritization strategy can be particularly powerful when combined with high-throughput functional assays like using MAVE model systems that are typically *in vitro*. Considering that both MAVE and proteogenomic datasets of tumor cohorts are both expanding quickly in the next few years^33,34^, the combined approaches can help effectively pinpoint functional mutations for mechanistic and clinical characterization. The prioritized mutations based on protein-level consequences may also guide the selection of targeted therapy to advance precision medicine.

## METHODS

### Proteogenomic datasets

The prospective CPTAC data were downloaded and processed as described in the Method section of the work of Elmas et al^35^. The overview table in **Figure 1A** of the dataset describes, for each cancer cohort, the sample size, female patient percentage, average cancer onset age, and tumor stage. Samples are normalized by their median absolute deviations (MAD), so that the MAD of all samples in the dataset is 1. Protein markers with high fractions (greater than 20%) of missing values are filtered out. For the corresponding RNA-seq data, we used the log2 normalization on the FPKM (fragments per kilobase of exon per million mapped fragments)-normalized RNA-seq counts and genes have no expression in at least 90% of the samples were filter out.

The proteomics data used for validation were downloaded from the NCI CPTAC portal. The dataset overview table in **Figure S1A** describe for each cancer cohort, the sample size, female patient percentage, average cancer onset age, and tumor stage. The validation data are processed in the same way as prospective data. The RNA-seq data sets of the three retrospective CPTAC cohorts were downloaded from the NCI CPTAC DCC portal. The RNA expression was measured in FPKM and was further normalized by log2(FPKM+1).

### pQTL and eQTL identification

For each cancer cohort, we identified pQTLs and eQTLs using the multiple linear regression model as implemented in the “limma” R package. We also corrected confounding factors including age, gender, ethnicity, and TMT batch. The false discovery rate (FDR) was corrected from the p-values with the Benjamini-Hochberg procedure. Somatic mutations are grouped at a gene level in the multiple regression model, similar to that implemented by our previously developed AeQTL tool^7^. Mutations separated are analyzed by their mechanisms of action, including nonsynonymous mutations as controls that likely do not affect expression, missense mutations, and truncating mutations including frameshift and in-frame indels, nonsense, splice site, and translation start site mutations. We focused on genes with three or more mutations in each cancer cohort and analyzed associations of mutations affecting *cis-*expression of the corresponding mRNA or protein products.

### spsQTL identification

We combined two statistical methods to identify spsQTLs. In the first method adopted from Battle et al.^4^, we compared the following two linear models using likelihood ratio test (LRT) with the “anova” function in R:

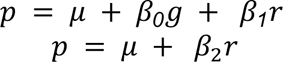

where *g* is the genotype, *r* represents RNA level, and p is the protein level. We filtered spQTLs that have an FDR less than 0.05 in LRT as candidate spsQTLs. In the complementary method adopted from Mirauta et al.^22^, we selected QTLs with a spQTL FDR less than 0.05 but an seQTL FDR greater than 0.05 as candidate spsQTLs. We then overlapped these two lists of candidate spsQTLs to identify the final list of spsQTLs for downstream analyses.

### Tumor-normal differential expression analysis

We conducted this analysis in the prospective CPTAC cohorts with paired tumor-adjacent tissure normal samples. For each cancer cohort, we paired the tumor and normal samples from the same patient and performed a differential protein/mRNA expression analysis to identify differentially expressed proteins with “limma” package. Demographic factors and batch effects, including age, ethnicity, race, and sequencing operator are adjusted in the multiple regression model.

## Supplementary Tables

**Supplementary Table 1. Significant spsQTLs identified by our statistical pipeline across six cancer types.**

**Supplementary Table 2. Test statistics between the three groups defined by (1) TP53 mutations with top 20% mRNA (left) or protein (right) expression in the prospective CPTAC cohorts, (2) the other TP53 mutations observed across all CPTAC samples, and (3) the rest of the assayed TP53 mutations from Kotler et al. using *TP53* functional scores form Kotler et al.**

## Supplementary Figures

**Supplementary Figure 1. Overview of the retrospective cohorts** (A) Summary of the retrospective CPTAC proteogenomic cohorts used for the discovery analyses, including cancer type abbreviation, data source, sample size of tumor (T) and normal (N) tissues, female percentage, average onset age in years, and tumor stage distribution. (B) Volcano plots showing seQTLs associations in the six cancer types (left) and volcano plots showing spQTLs associations (right), where each dot denotes a gene-cancer pair included in the analysis. Top associated genes were further labeled. FC: log fold change. FDR: false discovery rate.

**Supplementary Figure 2. spsQTLs with strong effects.** (A) Examples of spsQTL whose effect sizes in mRNA level and protein level are in different direction. For each gene, the plot on the left shows the corresponding mRNA levels of mutation carriers vs. non-carriers in FPKM, and the plot on the right shows protein level comparison in log ratio (MS TMT measurements) in the respective cancer type labeled on top of each of the violin plots. The labeled mutations are the three mutations whose carriers show the highest absolute expression differences of the mutated gene product compared to the non-carriers. (B) Examples of spsQTL with a protein logFC and mRNA logFC ratio greater than 15

**Supplementary Figure 3. Overlapped of significant QTLs in cross-tumor analysis and matched tumor-normal analysis projected onto pQTL volcano plots based on cross-tumor analyses.** The plots were made separately for (A) discordant spsQTLs, and (B) concordant eQTL/pQTLs.

**Supplementary Figure 4. Example lolliplots showing mutations for two genes that were identified as spsQTLs, including YLPM1 and ESRP1.** The number on each disc denotes the number of mutations in that position and the color of the disc represents the mutation type.

## DATA AND SOFTWARE AVAILABILITY

### Data Availability

Data for CPTAC cohorts can be found on CPTAC data portal: https://cptac-data-portal.georgetown.edu/cptacPublic/. Data for TP53 MAVE assays can be downloaded from the Supplementary Information from Kotler et al^21^.

### Code Availability

The source code used for all analyses in this article is available at https://github.com/Huang-lab/pQTL.

## ACKNOWLEDGEMENTS

The authors wish to acknowledge CPTAC and its participating patients and families that generously contributed the data. This work was supported by NIH NIGMS R35GM138113, ACS RSG-22-115-01-DMC, and Mount Sinai funds to KH.

## CONTRIBUTIONS

K.H. conceived the research; Y.L and K.H. designed the analyses. Y.L. developed the software and conducted the bioinformatics analyses, A.E. curated and preprocessed the datasets. Y.L. and K.H. wrote the manuscript. K.H. supervised the study. All authors read, edited, and approved the manuscript.

